# A mysterious 80 nm amoeba virus with a near-complete “ORFan genome” challenges the classification of DNA viruses

**DOI:** 10.1101/2020.01.28.923185

**Authors:** Paulo V. M. Boratto, Graziele P. Oliveira, Talita B. Machado, Ana Cláudia S. P. Andrade, Jean-Pierre Baudoin, Thomas Klose, Frederik Schulz, Saïd Azza, Philippe Decloquement, Eric Chabrière, Philippe Colson, Anthony Levasseur, Bernard La Scola, Jônatas S. Abrahão

## Abstract

Here we report the discovery of Yaravirus, a new lineage of amoebal virus with a puzzling origin and phylogeny. Yaravirus presents 80 nm-sized particles and a 44,924 bp dsDNA genome encoding for 74 predicted proteins. More than 90% (68) of Yaravirus predicted genes have never been described before, representing ORFans. Only six genes had distant homologs in public databases: an exonuclease/recombinase, a packaging-ATPase, a bifunctional DNA primase/polymerase and three hypothetical proteins. Furthermore, we were not able to retrieve viral genomes closely related to Yaravirus in 8,535 publicly available metagenomes spanning diverse habitats around the globe. The Yaravirus genome also contained six types of tRNAs that did not match commonly used codons. Proteomics revealed that Yaravirus particles contain 26 viral proteins, one of which potentially representing a novel capsid protein with no significant homology with NCLDV major capsid proteins but with a predicted double-jelly roll domain. Yaravirus expands our knowledge of the diversity of DNA viruses. The phylogenetic distance between Yaravirus and all other viruses highlights our still preliminary assessment of the genomic diversity of eukaryotic viruses, reinforcing the need for the isolation of new viruses of protists.

**Significance statement:** Most of the known viruses of amoeba have been seen to share many features that eventually prompted authors to classify them into common evolutionary groups. Here we describe Yaravirus, an entity that could represent either the first isolated virus of *Acanthamoeba* spp. out of the group of NCLDVs or, in alternative evolutive scenario, it is a distant and extremely reduced virus of this group. Contrary to what is observed in other isolated viruses of amoeba, Yaravirus is not represented by a large/giant particle and a complex genome, but at the same time carries an important number of previously undescribed genes, including one encoding a novel major capsid protein. Metagenomic approaches also testified for the rarity of Yaravirus in the environment.

## Introduction

Viral evolution and classification have been subject of an intense debate, especially after the discovery of giant viruses that infect protists (1–4). These viruses are predominantly characterized by the large size of their virions and genomes encoding hundreds to thousands of proteins, of which a large proportion currently remains without homologs in public sequence databases (5–9). These coding sequences are commonly referred as ORFans, and due to the lack of phylogenetic information, their origin and function still represent a mystery (10–13). Strikingly, the increasing number of available viral genomes demonstrate that there is a huge set and great diversity of genes without homologs in current databases, which need to be further explored (10). Importantly, many amoebal virus ORFan genes have already been proven to be functional, being expressed and encoding for components of the viral particles (6, 14). However, the large set of ORFans makes it difficult to predict the biology of viruses discovered through cultivation-independent methods, such as metagenomics analysis, reinforcing the need for the complementary isolation and experimental characterization of new viruses.

All currently known isolated amoebal viruses are related to nucleocytoplasmic large DNA viruses (NCLDVs) (15). This group comprises families of eukaryotic viruses (*Poxviridae, Asfarviridae, Iridoviridae, Ascoviridae, Phycodnaviridae, Marseilleviridae* and *Mimiviridae*) as well as other amoebal virus lineages including pithoviruses, pandoraviruses, molliviruses, medusaviruses, pacmanviruses, faustoviruses, klosneuviruses and others. NCDLVs have dsDNA genomes and were proposed to share a monophyletic origin based on criteria that include the sharing of a set of ancestral vertically inherited genes (16, 17). From this handful set of genes, a core gene cluster is found to be present in almost all members of the NCLDVs, being composed by five distinct genes, namely a DNA polymerase family B, a primase-helicase, a packaging ATPase, a transcription factor and a major capsid protein (MCP) for which the double jelly-roll (DJR) fold constitutes the main protein architectural class (18, 19). Recently, the International Committee on Taxonomy of Viruses (ICTV) brought forward a proposal for megataxonomy of viruses. The DJR major capsid protein (MCP) supermodule of DNA viruses is present in NCLDVs and other icosahedral viruses that infect prokaryotes and eukaryotes. In addition to the signature DJR-MCPs, the majority of these viruses also encode for additional single jelly roll minor capsid proteins (e.g., penton proteins) and genome packaging ATPases of the FtsK-HerA superfamily. According to this proposal, the evolutionary conservation of the three genes of the morphogenetic module in the DJR-MCP supermodule, to the exclusion of all other viruses, justifies the establishment of a realm named *Dividnaviria*. Although some NCLDVs, as pandoraviruses, seem to have lost the DJR-MCP gene, their genome present a large set of genes that support their classification into NCLDVs (and *Dividnaviria*, consequently).

Here we describe the discovery of Yaravirus, an amoeba virus with a puzzling origin and phylogeny. Yaravirus does not present all the three hallmark genes of *Dividnaviria*, lacking for an identifiable sequence of single jelly roll minor capsid protein. In addition, Yaravirus has an enigmatic genome whose near full set of genes are ORFans. Yaravirus can represent either the first isolated virus of *Acanthamoeba* spp. out of the group of NCLDVs. This virus has particles with a canonical size of 80 nm, escaping the concept of large and giant viruses. Thus, Yaravirus expands our knowledge about viral diversity and challenges the current classification of DNA viruses.

## Results

### Yaravirus isolation and replication cycle

A prospecting study was conducted by collecting samples of muddy water from creeks of an artificial urban lake called Pampulha, located at the city of Belo Horizonte, Brazil. Here, by using a protocol of direct inoculation of water samples on cultures of *Acanthamoeba castellanii* (Neff strain, ATCC 30010), we have managed to isolate a new amoebal virus that we named Yaravirus brasiliensis, as a tribute to an important character (Yara, the mother of waters) of the mythological stories of the Tupi-Guarani indigenous tribes (20). Negative staining revealed the presence of small icosahedral particles on the supernatant of infected amoebal cells, measuring about 80 nm diameter (Fig. 1a). Cryo-electron microscopy images of purified particles suggest that Yaravirus particles present two capsid shells, as previously described for Faustovirus (21).

**Figure 1.**
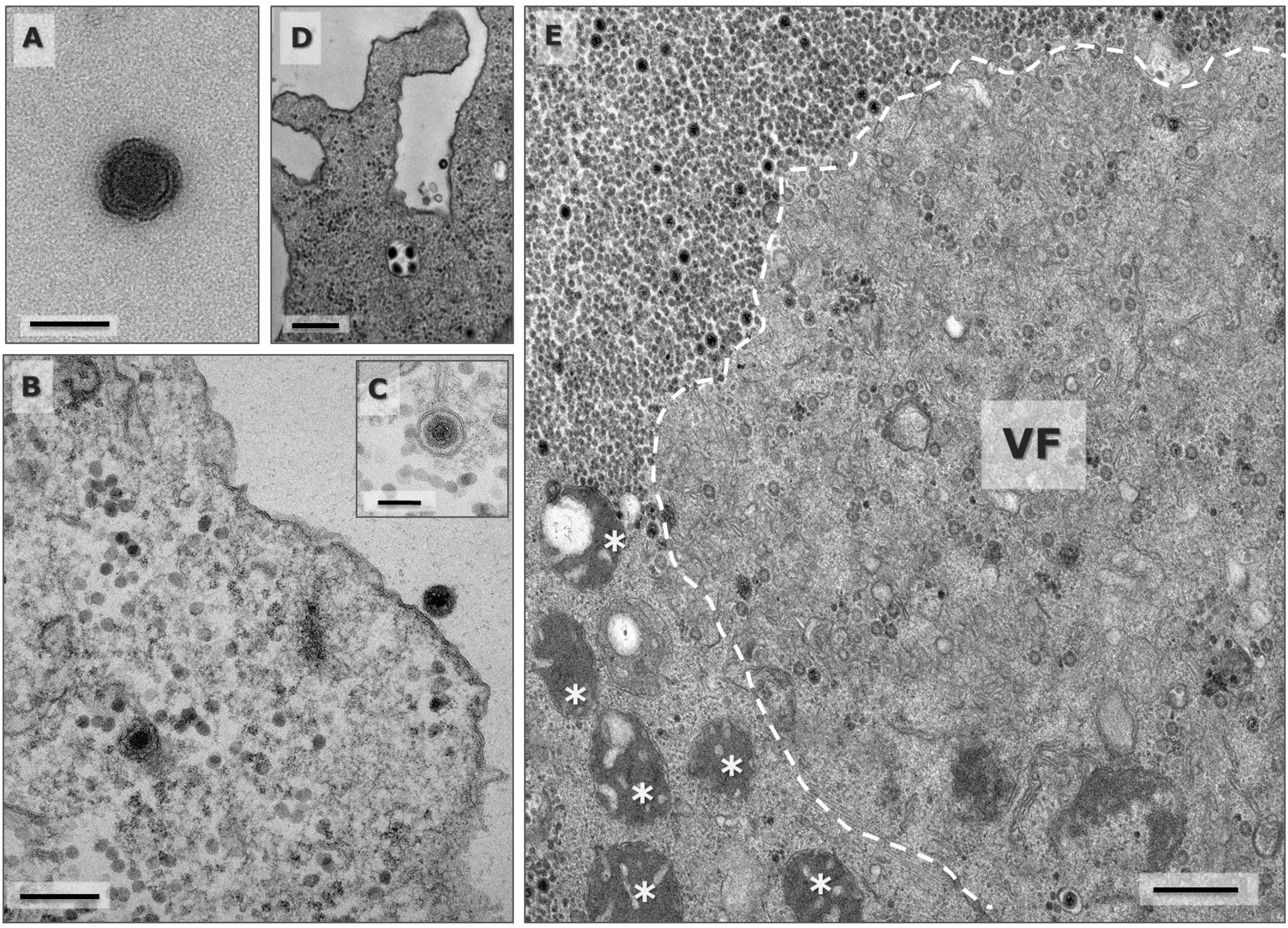
Yaravirus particle and the beginning of the viral cycle. **a** Negative staining of an isolated Yaravirus virion. Scale bar 100nm. **b** Transmission electron microscopy (TEM) representing the beginning of the viral cycle, in which one particle is associated to the host cell membrane and the second one was already incorporated by the amoeba inside an endocytic vesicle. Scale bar 200nm. **c** Detailed image of an incorporated Yaravirus particle in the interior of an endocytic vesicle. Scale bar 100nm. **d** Viral uptake by the amoeba may occur individually but also in groups of particles, as observed in the micrograph. Scale bar 250nm. **e** The viral factory completely develops occupying the nuclear region and recruiting mitochondria around it. Two different regions can be distinct: an electron-lucent region where the virions are assembled as empty shells and a second region formed by several electron-dense points where the genome is packaged inside the particles. Scale bar 500nm.

At the beginning of infection in *A. castellanii*, Yaravirus particles are found attached to the outside part of the amoebal plasma membrane, suggesting the participation of a host receptor in order to internalize the virions (Fig. 1b). The replication cycle is then followed by the incorporation of individual or grouped Yaravirus particles inside endocytic vesicles, which in a later stage of infection are found next to a region occupied by the nucleus (Fig. 1b-d). The viral factory then takes place and completely develops into its mature form, replacing the region formerly occupied by the cell nucleus and recruiting mitochondria around its boundaries, likely to optimize the availability of energy to construct the virions (Fig. 1e). The step corresponding to viral morphogenesis happens similarly as how it is observed for other viruses of amoeba. Firstly, it starts by the appearance of small crescents in the electron-lucent region of the factory (Fig. 1e, Fig. 2a). Next, step by step the virions gain an icosahedral symmetry by the sequential addition of more than one layer of protein or membranous components around its structure (Fig. 2a). The constructed virions, with a capsid still empty, start then to migrate to the periphery of the viral factory where there is the accumulation of corpuscular electron-dense material (Fig. 1e, Fig. 2b-c). These enucleations are scattered throughout the periphery of the infected cell and seem to represent different regions or morphogenesis points where the final step for Yaravirus maturation occurs. In these regions the capsid of Yaravirus is filled with electron-dense material and the virus is finally ready to be released (Fig. 2c – red arrow). Sometimes it is also possible to observe several particles of Yaravirus being packed in the interior of vesicle-like structures, suggesting a potential release by exocytosis, as observed for other viruses of amoeba (22, 23) (Fig. 2d). Most of the viral shedding, however, is still represented by lysis of the amoebal cell, followed by the release of Yaravirus particles, which later reach the supernatant of the infected culture, or sometimes might get attached to the debris of the cellular membranes (Fig. 2e). We have also evaluated Yaravirus replication by concomitantly investigating the decrease of the host cell numbers allied with the increase of viral genome during infection. Interestingly, during the first hours of infection, the *A. castellanii* cultures seem to progressively grow until 24 h.p.i, showing a fastidious character for Yaravirus replication (Fig. 2f). The cells then start to suffer lysis induced by the virus only after the 72 h.p.i (Fig. 2f). On the same level, from 96 h.p.i to 7 days post infection there is no change on the detection levels of Yaravirus genome and the lysis seems to stop, and the remaining trophozoites turn to cysts (data not shown).

**Figure 2.**
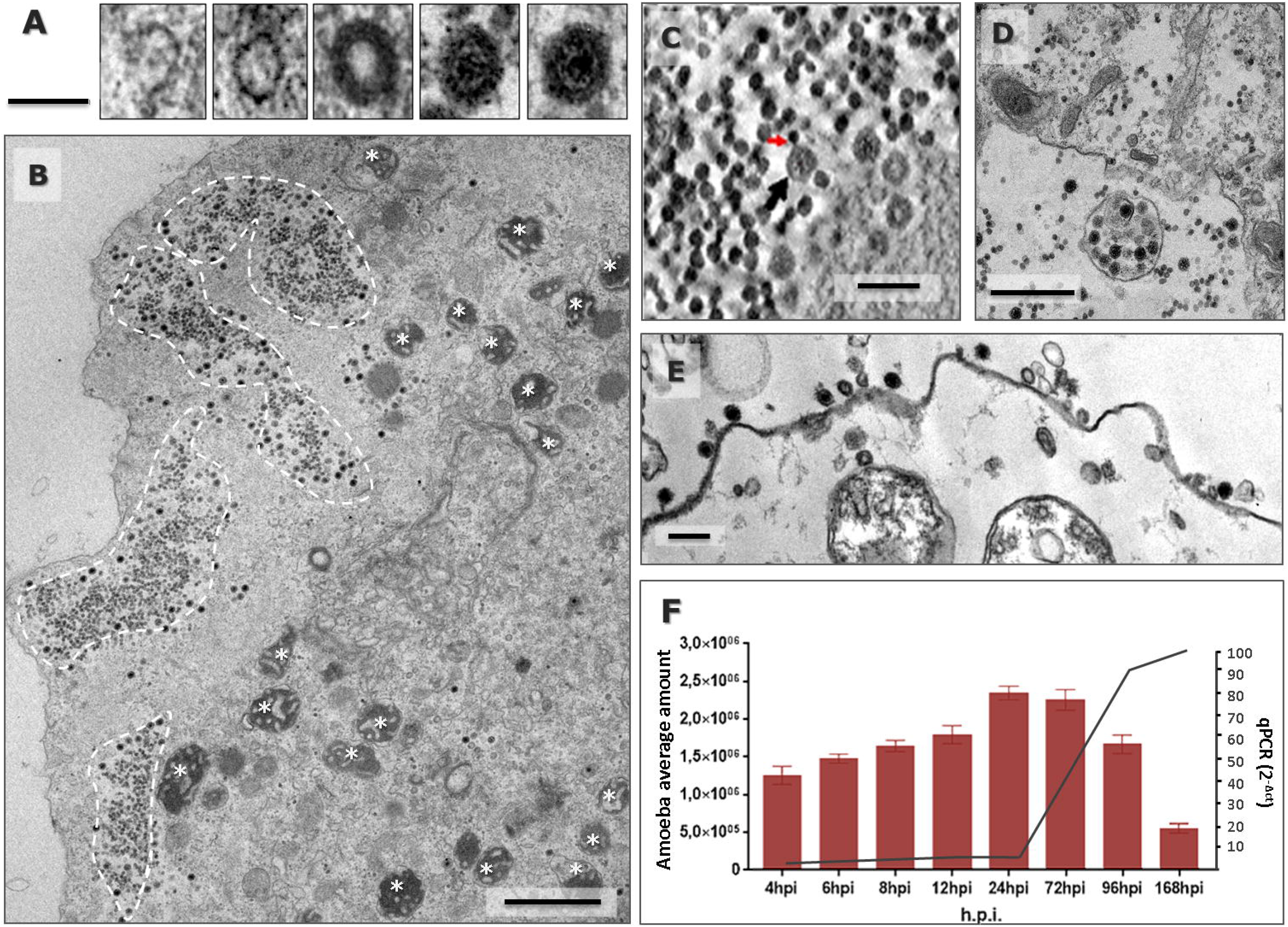
Yaravirus morphogenesis and release. **a** The virions are assembled by the addition of more than one layer of protein or membranous components around its structure. Scale bar 70nm. **b** The particles then start to migrate to the periphery of the cell where there is the presence of several electron-dense points that function as morphogenetic structures to package the DNA inside the Yaravirus particles (regions inside dashed lines). Scale bar 1000nm. **c** Detailed image of the morphogenetic regions where the DNA (red arrow) is incorporated inside the Yaravirus virion (black arrow). Scale bar 150nm. **d** Sometimes, the final step of viral replication is marked by the particles being packaged inside vesicle-like structures, suggesting a potential release by exocytosis. Scale bar 500nm. **e** Most of the particles, however, are released by cellular lysis and have a high affinity to the membranes of cellular debris. Scale bar 150nm. **f** Graph comparing concomitantly the decrease of host cell numbers (red bars) with the increase of Yaravirus genome during the infection (black line). Replication of viral genome was measure by qPCR and calculated by delta-delta Ct.

### Genome

Sequencing of the Yaravirus genome has shown the presence of a double-stranded DNA molecule with a length of 44,924 bp and harboring a total of 74 predicted genes (Fig. 3a). Despite a smaller genome than other viruses of amoeba, Yaravirus encodes for six tRNA genes: tRNA-Ser (gct), tRNA-Ser (tga), tRNA-Cys (gca), tRNA-Asn (gtt), tRNA-His (gtg) and tRNA-Ile (aat) (Fig. 3a and c). All of them are co-located on an intergenic region between genes 29 and 30 (Fig. 3a and c). In contrast to tRNA genes in tupanviruses, we did not observe a correlation between the Yaravirus tRNA isoacceptors and the codons most frequently used by the virus or its *A. castellanii* host. The genome has a GC content of 57.9%, which is one of the highest found in any amoebal virus discovered to date. When analyzed gene-by-gene, Yaravirus has a spectrum of GC content that varies between 46% and 65%. The analysis of the intergenic regions of the genome (46% GC content) did not reveal any enriched sequence motifs that might indicate a conserved promoter, opposed to what is observed in many other NCLDV members (24).

**Figure 3.**
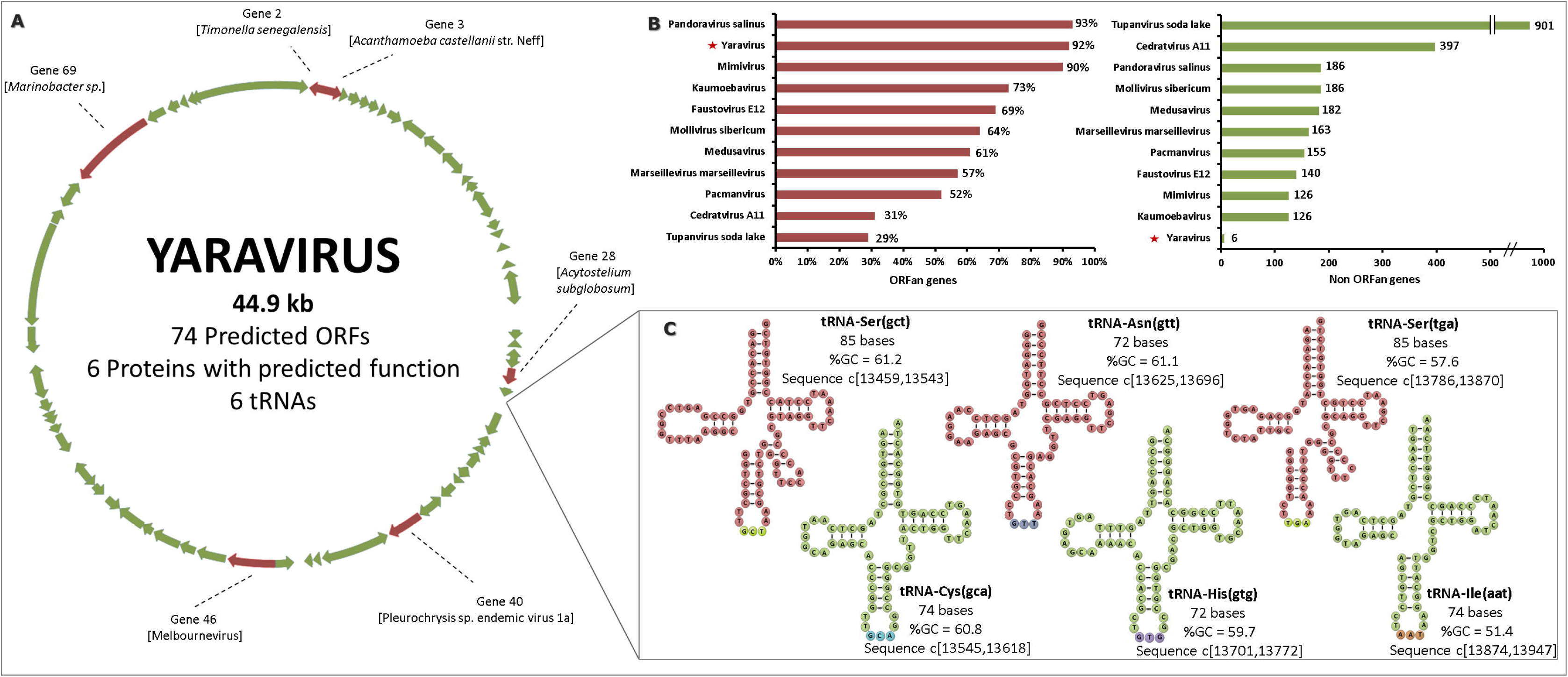
Yaravirus genome features. **a** Circular representation of Yaravirus genome highlighting the only six genes (red arrows) which have similarity with aminoacid sequences of other organisms in current databases. Genes with no matches on the databases are represented as green arrows. **b** The percentage of ORFan genes among the complete genome of different viruses of amoeba is represented by the graph with red scale bars. The graph with greenish scale bars represents the absolute number of genes with homologues in databases (non ORFan genes) for each of the same amoebal viruses previously analyzed. **c** All the six Yaravirus predicted tRNAs, as well as their corresponding sequences, are pictured with information about their anticodon (in parenthesis), their nucleotide length, the % of GC content and the position in the intergenic regions of genes 29 and 30.

By considering only the portions of genome related to coding regions, Yaravirus also has a similar coding capacity as observed in other viruses when their genome was first annotated, approximately 90%.

Surprisingly, Yaravirus genome annotation showed that none of its genes matched with sequences of known organisms when we compared them at the nucleotide level. When we looked for homology at the aminoacid levels, we found that only two predicted proteins had hits in the Pfam-A database and in total six had distant matches in the nr database. Complimentary prediction of three-dimensional structures of these proteins indicated a potential function of 4 more genes (see the proteomics analysis). Taken together, more than 90% (68) of the Yaravirus predicted genes are ORFans, something not observed for amoebal viruses since the discovery of the pandoraviruses (even after using a more relaxed criteria, BLASTp, e-value < 10^−3^) (Fig. 3b) (25). The six genes whose product has some homology with known protein sequences (Fig. 3b) (Table 1) are homologous to fragments of proteins predicted to have different functions, such as an exonuclease/recombinase bacterial protein (gene 2; best hit: *Timonella senegalensis*), a hypothetical protein (gene 3; best hit: *A. castellanii*), a hypothetical protein (gene 28; best hit: *Acytostelium subglobosum LB1* - a dictyostelid), a packaging ATPase (gene 40; best hit: Pleurochrysis endemic virus), a conserved hypothetical protein (gene 46; best hit: Melbournevirus, a marseillevirus strain) and a bifunctional DNA primase/polymerase (gene 69; best hit: *Marinobacter sp* – *Alpha* proteobacteria) (Table 1).

**Table 1.** Yaravirus genes with similarity on current databases and their best-hits

| **Yaravirus**<br>**Gene ID** | **Best**<br>**Hit** | **Total**<br>**score** | **Query**<br>**cover** | **E-value** | **Identity** | **Annotation of**<br>**best blastp hit** |
| --- | --- | --- | --- | --- | --- | --- |
| 2 | Timonella senegalensis WP_019148817.1 | 67.0 | 87% | 1e-09 | 27.88% | exonuclease/recombinase |
| 3 | Acanthamoeba castellanii XP_004339080.1 | 51.6 | 80% | 1e-06 | 44.12% | hypothetical protein |
| 28 | Acytostelium subglobosum LB1<br>XP_012747655.1 | 57.8 | 85% | 4e-07 | 27.59% | hypothetical protein |
| 40 | Pleurochrysis sp. endemic virus 1a<br>AUD57256.1 | 121 | 66% | 4e-28 | 33.05% | virion packaging ATPase |
| 46 | Melbournevirus YP_009094634.1 | 181 | 88% | 5e-47 | 32.63% | hypothetical protein conserved in<br>marseilleviruses |
| 69 | Marinobacter sp. MAB50943.1 | 106 | 35% | 8e-20 | 25.24% | bifunctional DNA primase/polymerase |

Phylogenetic analyses were then performed for those different genes (genes 02, 03, 28, 40, 46 and 69) after aligning them with protein sequences of similar function belonging to different members of the virosphere and to organisms of the three cellular domains of life. Given the lack of representatives on already known databases other than their best-hits, sequences corresponding to the genes 03 and 28 (both hypothetical proteins) didn’t have enough genetic information to be included in a phylogenetic analysis. For analysis corresponding to the gene 02 (exonuclease/recombinase), three major groups were observed to construct the morphology of the tree. Yaravirus was observed to be placed in one of those branches, clustering with some members of Eukarya, specifically with stony coral and insects (Fig. 4). Analyses of gene 40 (virion packing ATPase) revealed that Yaravirus clustered in a polyphyletic branch, with members belonging to *Mimiviridae* family, bacteria (although many of these sequences seem to represent misclassified NCLDVs from metagenome-assembled genomes) and Pleurochrysis sp. endemic virus 1a and 2. For the phylogenetic analyses corresponding to gene 46 (hypothetical protein conserved in Marseillevirus), Yaravirus clustered with Marseillevirus strains. For the last tree, representing analysis for gene 69 (bifunctional DNA primase/polymerase), we have observed that Yaravirus shares a cluster with members of eukaryotes corresponding to the Streblomastix and Phytophthora groups. However, it should be noted that in a previous study the authors detected sequences of mimivirus genes among the Phytophthora parasitic strain INRA-310 genome (26). After all those analyses, it is important to note that although Yaravirus has some genes with representatives in the genome of other organisms, their homology with orthologs is very low (25.24 to 44.12%), highlighting that Yaravirus genome content is essentially novel among the other members of the virosphere (Table 1).

**Figure 4.**
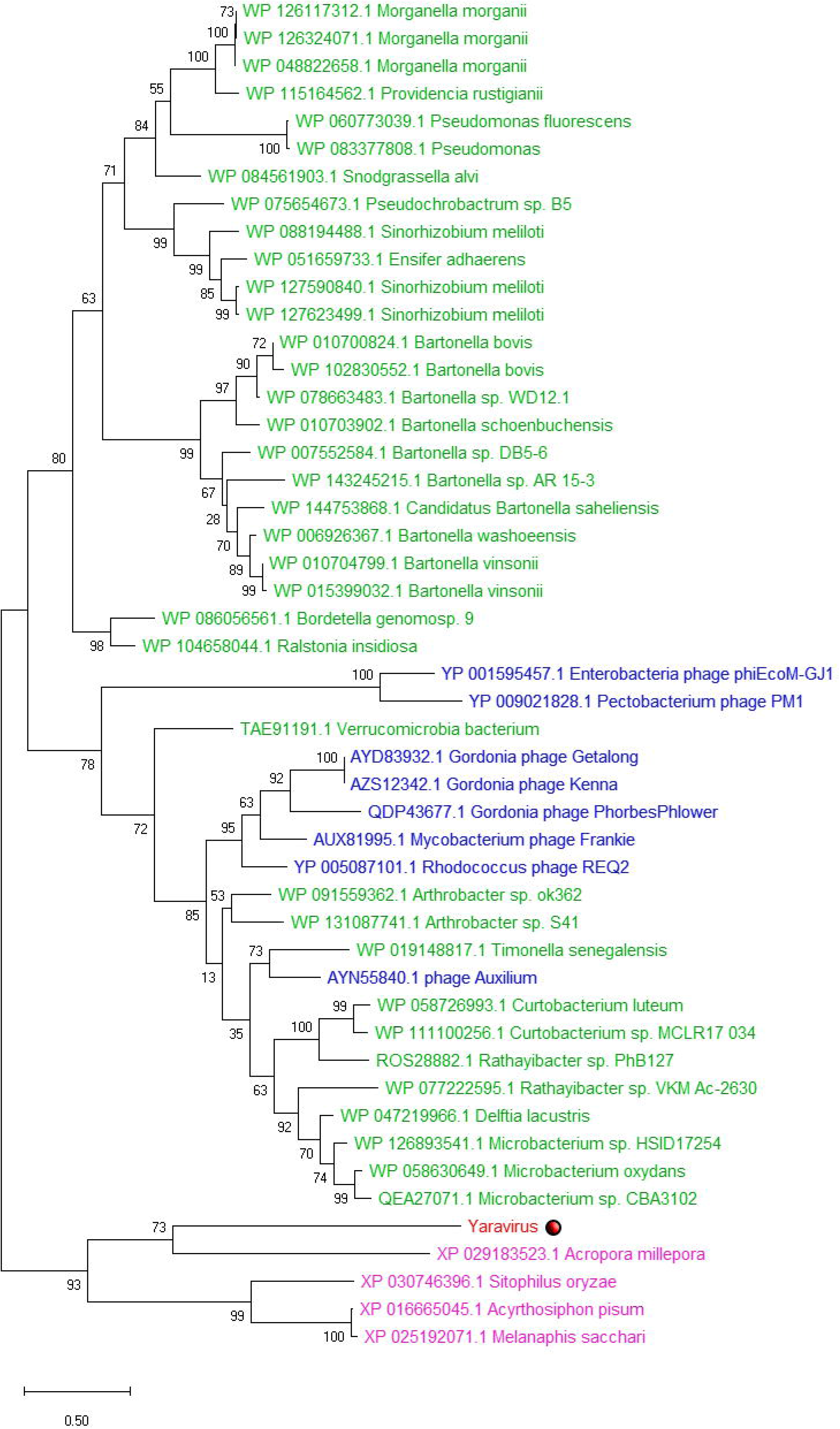
Phylogenetic tree of the Yaravirus gene corresponding to a probable exonuclease/recombinase, presented in the best-hit *Timonella senegalensis*. Similar genes incorporated in the genome of bacteria (green), eukarya (pink) and other viruses (blue) are also represented.

In order to detect sequences related to Yaravirus we surveyed 8,535 publicly available metagenomes in the IMG/M database that have been generated from samples from diverse habitats across our planet (27). We discovered distant homologs of the Yaravirus ATPase (NCVOG0249) with an amino acid homology of up to 33.9% in the metagenomic data, while the closest homolog in the NCBI nr database was that of Pleurochrysis sp. endemic virus 1a with 33.1%. In a phylogenetic tree of the viral ATPase the Yaravirus branched within the *Mimiviridae* as part of a highly supported clade made up by its distant metagenomic relatives and three viruses whose genomes were deposited in NCBI Genbank and named Pleurochrysis sp. endemic virus 1a, 1b and 2 (Fig. 5). In contrast to known members of the *Mimiviridae*, viral contigs and viral genomes in this clade featured a high GC content with up to 62%. We also searched for proteins similar to the Yaravirus putative MCP but were not able to retrieve closely related sequences in the metagenomic data.

**Figure 5.**
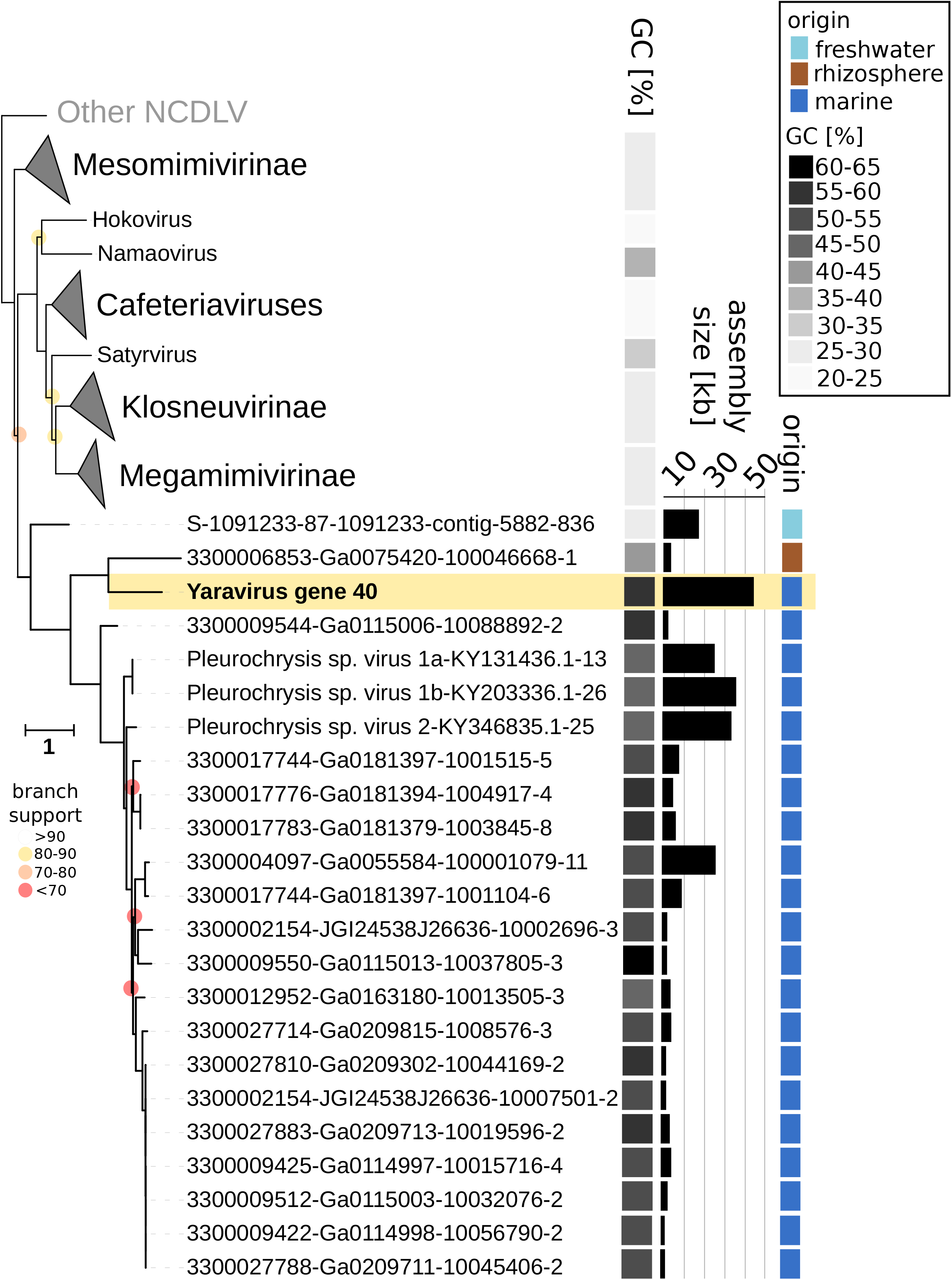
Phylogenetic position of Yaravirus and related viral sequences in the *Mimiviridae* based on the viral ATPase (NCVOG0249). The Yaravirus ATPase is highlighted in yellow. Branch support is indicated as colored circles for support values of 90 or below. The tree is rooted at the Poxviruses. Scale bar indicates substitutions per site. GC content of viral genomes and contigs containing NCVOG0249 is shown together with the average GC content of collapsed clades. In addition, environmental origin and assembly sizes of Yaravirus and related viral contigs and genomes are shown. The accession numbers (IMG/M)(REF) of metagenomic sequences are indicated as the numbers before the first dash in the sequence name.

### Yaravirus proteomics

As aforementioned, most Yaravirus proteins had no detectable homologs in public databases and, from a first perspective, the virus did not encode for capsid proteins. This peculiarity prompted us to have a closer look at the proteins responsible to form the mature particles of Yaravirus. Proteomics revealed a total of 26 viral proteins present in purified particles. We then analyzed the predicted three-dimensional structures of those 26 proteins, by using two platforms for domain comparison, the Phyre2 and the Swiss-model tools (28–33). Remarkably, only 4 sequences (genes 11, 12, 41 and 46) were observed to have structural features similar to known proteins (Table 2). That means that almost 90% of its virion proteome consists of ORFans. It is important to mention that the same approach (in silico structure prediction) has been used in parallel to evaluate all of the 74 predicted genes on the genome of Yaravirus, resulting in the discovery of two more genes with structural resemblance with other proteins in public databases (gene 02 and 70) (Table 2).

**Table 2.** Annotation of Yaravirus proteins based on the predicted tridimensional structure of the proteins coded by the virus.

| Gene | Hits with Phyre | Sequence cover | Confidence | Swiss-Model |
| --- | --- | --- | --- | --- |
| 2 | Endonuclease_Exonuclease | 76% | 100% | Exonuclease |
| **11** | **Signal protein TNF-like** | **44%** | **93%** | **Complement C1q-like protein** |
| **12** | **Signal protein TNF-like** | **55-62%** | **96%** | **Complement C1q-like protein** |
| 40 | ATPase | 57% | 99.6% | Replicative DNA helicase |
|  | DNA translocase | 46% | 98.7% |  |
| **41** | **Capsid** | **65%** | **97%** | **Capsid** |
| **46** | **Secreted protein PcsB** | **75%** | **99.2%** | **Tail needle protein** |
|  | **Capsid** | **26-49%** | **98%** |  |
| 70 | Endonuclease holliday junction resolvase | 77% | 95% | Holliday junction resolvase |

Note: the bold-type genes represent proteins observed on the viral proteomics

Proteomics data revealed that the most abundant proteins in the viral particles corresponded to genes 41, 46 and 51 (from most to less abundant) (Table 2). While for the third highest expressed protein we were not able to find any structural candidates with known biological function, for sequences represented by the genes 41 and 46 we observed fragments of protein resembling the three-dimensional structure of the capsid of other viruses (Table 2). With a confidence of 97%, a relevant portion (65%) of the gene 41 was found to have a structural convergence with the double-jelly roll domain of the MCP of the Paramecium bursaria Chlorella virus type 1. Gene 46 encoded predicted protein was found to have structural convergence with bacterial secreted protein pcsB and tail needle protein, a portion composed of a long alpha-helix. The function of protein encoded by gene 46 remains to be investigated. Therefore, we were not able to find convincingly any minor capsid protein. It is also important to note that sequences represented by the gene 46 are the same described earlier to be highly conserved (although with unknown function) in marseilleviruses and in medusaviruses (Table 1). Finally, the last two proteins observed in the proteomics which had structural deposits in public databases are represented by the genes 11 and 12, both expressing predicted complement C1-qlike proteins.

## Discussion

In the last years, amoebal large and giant viruses have frequently been found around the world (5–9, 20, 22, 25, 34–37). Here, we describe Yaravirus brasiliensis, an 80 nm-sized virus with a genome containing a notable proportion of genes (~90%) that have never been observed before. Using standard protocols, our very first genetic analysis was unable to find any recognizable sequences of capsid or other classical viral genes in Yaravirus. This is a relevant feature to highlight the importance of studies related to the isolation of new viral samples, as by following the current metagenomic protocols for viral detection, Yaravirus would not even be recognized as a viral agent (38, 39). According to our knowledge, Yaravirus represents the first virus isolated in *Acanthamoeba* spp. that is potentially not part of the complex group of NCLDVs. Several characteristics unite previously discovered amoebal viruses: large-sized virions, genomes coding for hundreds to thousands of genes and presumably a monophyletic origin that is reflected in the presence of a set of about 20 most likely vertically inherited genes (16, 17). None of these features are present in Yaravirus, and that makes it potentially the first isolate of a novel *bona fide* group of amoebal virus. Of course, we cannot exclude the possibility that Yaravirus may represent a reduced NCLDV, presenting highly divergent or even absent NCLDV hallmark proteins. Recently, a similar case was described for three small crustacean viruses. However, despite their reduced genome when compared to other members of the NCLDV, an important number of hallmark genes were shared with this group, differently as observed for Yaravirus (40). In this scenario, not less exciting, Yaravirus would represent the to-date smallest member of the NCLDVs, both in particle and genome size. The presence of six copies of tRNAs in Yaravirus also impresses when analyzed by the perspective of a selective pressure forcing to maintain these genes in such a small genome, when compared to larger viruses of amoeba. Even more interestingly, none of the isoacceptors related to the Yaravirus tRNAs corresponds to codons of amino acids abundantly used by the virus or the amoeba. Considering the fastidious infection cycle of Yaravirus in *Acanthamoeba*, it is conceivable that in nature a different organism might act as the preferred host of Yaravirus.

Most members of the to date isolated giant viruses of amoeba show a capsid specially composed by copies of an MCP related to the D13L of *Vaccinia virus* (14, 41). Pandoraviruses are an exception as they seem to lack a protein shell to protect their genomes (25). Interestingly, even some of the amoeba hosts of these viruses may carry copies of MCP genes, suggesting possible horizontal gene transfer between virus and protist host (42, 43). Even though the Yaravirus capsid does not seem to be homologous to the NCLDV MCP, one of its most abundant proteins features the same architectural double-jelly roll observed in the MCP of giant viruses. This highlights how proteins with completely undescribed sequences might be shaped by evolutionary convergence to play important biological functions (44). Taken together, we can conclude that Yaravirus represents a new lineage of viruses isolated from *A. castellanii* cells. The amount of unknown proteins composing the Yaravirus particles reflects the variability existing in the viral world and how much potential of new viral genomes are still to be discovered.

## Methods

### Origin of samples and viral isolation

In 2017, searching to isolate novel variants of virus-infecting amoebas, we have collected samples of muddy water from a creek of the Pampulha lake, an artificial lagoon located at the city of Belo Horizonte, Brazil (19◦51 0.60S and 43◦58 18.90W). As soon as they were collected, the samples were quickly taken to our lab and stored at 4°C until they were further processed. Following the protocol, 4×10^4^ amoebas of the *Acanthamoeba castellanii* Neff strain (ATCC 30010) were seeded in each well of a 96-well plate, inoculating to each one a volume of around 100uL of the collected samples, originally diluted 1:10 in PBS buffer. The plates were then incubated for 7 days at 32°C and observed daily for the appearance of cytopathic effect, what may indicate a probable viral infection. All the content from the wells was then collected and submitted to three processes of freezing and thawing and analysis of the possible isolates by negative staining technique. By the end, the collected content was submitted to another two blind passages in fresh cultures of amoeba, but this time, in 25 cm^2^ Nunc™ Cell Culture Treated Flasks with Filter Caps (Thermo Fisher Scientific, USA) containing around 1 million amoebal cells. After viral isolation, all the following experiments were made by infecting *Acanthamoeba castellanii* cells in a low multiplicity of infection (MOI), given the Yaravirus fastidious replication cycle.

### Transmission electron microscopy (TEM), TEM tomography, cryo-electron microscopy

For resin embedding and transmission electron microscopy (TEM) *Acanthamoeba castellanii* cells infected with Yaravirus were fixed at 20 hours post-infection with 2.5 % glutaraldehyde in 0.1M sodium cacodylate buffer. Cells were washed three times with a solution of 0.2M saccharose in 0.1M sodium cacodylate. Cells were post-fixed for 1h with 1% OsO4 diluted in 0.2M Potassium hexa-cyanoferrate (III) / 0.1M sodium cacodylate. After washes with distilled water, cells were gradually dehydrated with ethanol by successive 10 min baths in 30, 50, 70, 96, 100 and 100 % ethanol. Substitution was achieved by successively placing the cells in 25, 50 and 75 % Epon solutions for 15 min. Cells were placed for 1 h in 100 % Epon solution and in fresh Epon 100 % over-night at room-temperature. Polymerization took place with cells in fresh 100 % Epon for 48 h at 60°. Ultra-thin 70 or 300 nm thick sections were cut with a UC7 ultramicrotome (Leica) and placed on HR25 300 Mesh Copper/Rhodium grids (TAAB, UK). Ultra-thin sections were contrasted according to Reynolds (45). Electron micrographs were obtained on a Tecnai G^20^ TEM operated at 200 keV equipped with a 4096◻×◻4096 pixels resolution Eagle camera (FEI)). For tomography, gold nanoparticles 10 nm in diameter (Ref. 741957; Sigma-Aldrich) were deposited on both faces of the sections prior to contrasting. Tomography tilt series were acquired on the G^20^ Cryo TEM (FEI) with the Explore 3D (FEI) software for tilt ranges of 110° with 1° increments. The mean applied defocus was −2 μm. The magnification ranged between 3500 and 29,000 with pixel sizes between 3.13 and 0.37 nm, respectively. The image size was 4096^2^ pixels. The tilt-series were aligned using ETomo from the IMOD software package (University of Colorado, USA) by cross-correlation (46). The tomograms were reconstructed using the weighted-back projection algorithm in ETomo from IMOD. The average thickness of the obtained tomograms was 268,40 ± 64 nm (*n* = 16). Fiji/ImageJ (NIH, USA) was used for making tomography movies (47).

For cryo-electron microscopy assays, the supernatant of infected cultures of *Acanthamoeba castellanii* was collected after 7 days post-infection and submitted to a first round of centrifugation, at 1500g for 10 min, looking to pellet the cell debris from the virus present on the supernatant. Next, the portion containing the Yaravirus was then submitted to a second round of centrifugation, and the virus was concentrated by ultracentrifugation at 100.000g for 2h. The following steps were previously described by Klose et al (21). Briefly, the μl of virus solution were placed on glow discharged C-Flat 2/2 grids (EMS) and plunge frozen into liquid ethane using a Gatan Cryoplunge 3. Samples were then imaged on a Talos F200C (ThermoFisher Scientific) equipped with a Ceta camera (ThermoFisher Scientific).

### Genome sequence and analysis

The Yaravirus genome was sequenced two times by using the Illumina Miseq platform (Illumina Inc., San Diego, CA, USA) with the paired-end application. The generated reads were then assembled *de novo* by using the software ABYSS and SPADES, with the resulting contigs ordered by the Python-based CONTIGuator.py software. After, gene predictions were made by using the GeneMarkS tool (48). The functional annotation for the Yaravirus predicted proteins was made through searches against the GenBank NCBI non-redundant protein sequence database (nr), considering homologous proteins only the sequences that presented an e-value < 1×10^−3^. For the qPCR assays, the increase in genome replication was assessed in cultures of *Acanthamoeba castellanii* cells infected by Yaravirus in different time points (H4, H6, H8, H12, H24, H72, H96 and H168), using primers which were constructed based on the sequence of the gene 69 of Yaravirus (primers: 5’TGCAGCAAGTCGGTCAAGAT3’ and 5’AACTTCCACATGCGAAACGC3’). Conditions used in the assay were previously described (49).

The aminoacid and codon usage data was compared to those presented by *Acanthamoeba castellanii* and by different strains of amoebal viruses. For this, the sequences were downloaded from the NCBI database and analyzed by using the software Artemis 18.0.3. The % of GC content and GC skew have also been analyzed by using the same software. Transfer RNA (tRNA) sequences were identified using the ARAGORN tool.

Phylogenetic analyses were performed for the six proteins of Yaravirus holding similarities with other organisms on the NCBI database (Table 1). By using the Clustal W tool in the Mega 10.0.5 software program, aminoacid sequences of these Yaravirus proteins were previously aligned with the corresponding sequences of representatives of the virosphere and from other cellular organisms belonging to the three Domains of Life. All the trees were constructed by using the maximum likelihood evolution method, with the JTT matrix-based model and a bootstrap of 1000 replicates (50).

### Yaravirus proteomics

In order to identify the proteins that make up Yaravirus particles, thirty 75cm2 cell culture flasks (Nunc, USA), containing 7×10^6^ *Acanthamoeba castellanii* cells/flask, were infected with the isolated virus and the cytopathic effect was followed up to 7 d.p.i. After severe amoebal lysis, the content was collected and submitted to a first round of centrifugation, at 1500g for 10 min, looking to pellet the cell debris from the virus present on the supernatant. Then, this viral portion was submitted to a second round of centrifugation, and the virus was concentrated by ultracentrifugation at 100,000g for 2h. To finish, viral pellet was then prepared for a 2D gel electrophoresis and analysis by MALD-TOF and LC-MS/MS as described before by Reteno and colleagues (51)

### Metagenomic survey

The Yaravirus ATPase (NCVOG0249) and the putative Major Capsid Protein (MCP) were used to query 8,535 publicly available metagenomes in the IMG/M database (27) using diamond blastp (v0.9.25.126, (52)). Resulting protein hits with more than 30% query and subject coverage and an E-value of at least 1e-5 were extracted from the metagenomic data. In parallel, hmmsearch (version 3.1b2, hmmer.org) was employed to identify and extract ATPases (NCVOG0249) and MCPs from 235 NCDLV reference genomes using specific hidden Markov models (https://bitbucket.org/berkeleylab/mtg-gv-exp/). Extracted proteins were then combined with the Yaravirus queries, aligned with MAFFT-linsi (v7.294b,(53)) (ATPase) and MAFFT (MCP) and the resulting amino acid alignments trimmed with trimal (v1.4, -gt 0.9, (54)). Phylogenetic trees were built using IQ-tree (v1.6.12, (55)) with LG+F+R5 (ATPase) and LG+F+R8 (MCP) based on the built-in model select feature (56) and 1000 ultrafast bootstrap replicates (57). The ATPase phylogenetic tree was visualized with iTol (v5, (58)).

## Data availability

Yaravirus genome and proteomics data will be available upon the manuscript publication.

## Acknowledgments

We thank our colleagues from IHU (Aix Marseille University) and from Laboratório de Vírus (Universidade Federal de Minas Gerais) for their assistance, specially Said Mougari, Issam Hasni, Lina Barrassi, Priscilla Jardot, Erna Kroon, Claudio Bonjardim, Paulo Ferreira, Giliane Trindade and Betania Drumond. In addition, we thank the Méditerranée Infection Foundation, Centro de Microscopia da UFMG, CNPq (Conselho Nacional de Desenvolvimento Científico e Tecnológico), CAPES (Coordenação de Aperfeiçoamento de Pessoal de Nível Superior) and FAPEMIG (Fundação de Amparo à Pesquisa do estado de Minas Gerais) for their financial support. J.A. is a CNPq researcher. B.L.S., J.A., P.C., P.V.M.B. and G.P.O. are members of a CAPES-COFECUB project.

